# Quantitative *in vivo* bioluminescence imaging of orthotopic patient-derived glioblastoma xenografts

**DOI:** 10.1101/2020.05.15.098640

**Authors:** Anna L. Koessinger, Dominik Koessinger, Katrina Stevenson, Catherine Cloix, Louise Mitchell, Anthony J. Chalmers, Jim C. Norman, Stephen W.G Tait

## Abstract

Despite extensive research, little progress has been made in glioblastoma therapy, owing in part to a lack of adequate preclinical *in vivo* models to study this disease. To mitigate this, primary patient-derived cell lines, which maintain their specific stem-like phenotypes, have replaced established glioblastoma cell lines. However, due to heterogenous tumour growth inherent in glioblastoma, the use of primary cells for orthotopic *in vivo* studies often requires large experimental group sizes. Therefore, when using intracranial patient-derived xenograft (PDX) approaches, it is advantageous to deploy imaging techniques to monitor tumour growth and allow stratification of mice. Here we show that stable expression of near-infrared fluorescent protein (iRFP) in patient-derived glioblastoma cells enables rapid direct non-invasive monitoring of tumour development without compromising tumour stemness or tumorigenicity. Moreover, as this approach does not depend on the use of agents like luciferin, which can cause variability due to changing bioavailability, it can be used for quantitative longitudinal monitoring of tumour growth. Notably, we show that this technique also allows quantitative assessment of tumour burden in highly invasive models spreading throughout the brain. Thus, iRFP transduction of primary patient-derived glioblastoma cells is a reliable, cost- and time-effective way to monitor heterogenous orthotopic PDX growth.

## Introduction

Glioblastoma is the most common malignant intrinsic brain tumour in adults ^1^. Current treatment regimens, consisting of surgical resection followed by radio- and chemotherapy, have limited efficacy in combating the disease, and this is reflected in a dismal prognosis for the patients ^2,3^. Improvement of therapeutic strategies is urgently needed to increase patient life expectancy and to reduce the disabling effects of the disease and its therapy. Development of novel therapies implicitly depends on reproducible and reliable models for preclinical studies *in vitro* and *in vivo*. However, experiments using orthotopic xenografts of established glioblastoma cell lines in immunocompromised mice to investigate effects of new therapies have many limitations. Most importantly, positive results observed in these preclinical studies have not been reproduced in subsequent clinical studies ^4,5^.

To better recapitulate human disease, the use of patient-derived glioblastoma stem-like cells (GSC) has gained increasing prominence in recent years ^6,7^. Several studies have demonstrated that patient-derived GSC cultured in serum-free neurobasal medium, supplemented with epidermal growth factor (EGF) and fibroblast growth factor (FGF), are more representative of primary glioblastoma than traditional, commercially-available glioblastoma cell lines ^8,9^. Nevertheless, it is not trivial to maintain the GSC population of surgical specimens using *in vitro* cell culture systems as stem cell growth varies due to plasticity, resulting in artificial selection of proliferating, rather than dormant GSC. Therefore, it is advantageous to study glioblastoma behaviour *in vivo,* as brain tissue provides the most appropriate microenvironment for maintenance of the stem signature of patient-derived xenografts (PDX) ^7^. However, due to their intrinsic heterogeneity, orthotopic PDXs display a much greater range of growth rates between animal subjects compared to established glioblastoma cell lines ^4,10^. Although this variability is a good reflection of the heterogeneity of the parental tumour, differences in growth characteristics, and therefore PDX size and the degree of dissemination at defined time points, lead to difficulties in quantifying and interpreting animal studies. Thus, large cohorts of animals are needed to achieve the necessary statistical power in these preclinical studies.

Currently, intra-vital bioluminescent imaging (BLI) with luciferase is commonly used to monitor xenograft growth ^11^. Yet, *in vivo* luciferase assays are limited to descriptive analysis since factors such as fluctuating substrate access to the target tissue result in variable luciferin bioavailability in tumours. Moreover, luciferase-based BLI does not allow for absolute quantification of the same tumour over time. Although magnetic resonance imaging (MRI) is ubiquitously used as clinical standard of care to diagnose brain tumours in humans, this technique has major drawbacks in preclinical settings. First of all, small animal MRIs are not widely available. Secondly, the procedure and its analysis are expensive as well as onerous for animals, requiring prolonged episodes of anaesthesia. Thirdly, MRIs often only measure indirect indicators of tumour pathology, such as focal brain oedema, which do not necessarily reflect the size and margins of the tumour itself.

To address these challenges, we have adapted a method of viral transduction to express iRFP, a protein with excitation and emission in the near-infrared spectrum, in patient-derived GSC, whilst maintaining their stem-like phenotype and growth characteristics. Using these cells, we find that imaging iRFP is a reliable, rapid and cost-effective method to monitor tumour bulk and infiltrative growth of orthotopic glioblastoma models *in vivo.*

## Materials and Methods

### Patient-derived GSC and Cell culture reagents

Patient-derived E2 and G7 glioblastoma stem-like cells (GSC), obtained from surgical resection specimens of anonymised patients as previously described ^12,13^, were kindly provided by Prof. Colin Watts. Serum-free medium for GSC was Advanced Dulbecco’s modified Eagle’s medium F12 (Thermo Fisher Scientific), supplemented with 2mM glutamine, 4μg/ml heparin (Sigma), 1% B27 (Thermo Fisher Scientific), 0.5% N2 (Thermo Fisher Scientific), 20ng/ml EGF and 10ng/ml FGF (Thermo Fisher Scientific). GSC were cultured in a 37°C incubator at 5% CO_2_ and, when grown as monolayers, on Matrigel™ (Corning) pre-coated plates. For all experiments GSC were used up to five passages after thawing. Cell lines were routinely tested for mycoplasma.

### iRFP vector and retroviral transduction

As described previously, plasmids encoding iRFP IRES puro had been inserted into a pBABE vector ^11^. For GSC transduction HEK293-FT cells (4×6^6^ in a 10 cm dish, cultured in high-glucose DMEM, complemented with 10% FCS and 2mM glutamine) were transfected using 4μg polyethylenimine (PEI, Polysciences Inc., Warrington, USA) per μg plasmid DNA. The retroviral transfer vector plasmid pBABE iRFP IRES puro (kindly provided by Andreas Hock ^11^), packaging plasmid HIV-gag-pol (Addgene 14887) and envelope plasmid pUVSVG (Addgene 8454) DNA were mixed in a 4:2:1 ratio. DNA/PEI mixtures were incubated at room temperature for 10 to 15 minutes, prior to application on HEK293-FTs. 24 and 48 hours later, virus containing supernatant was harvested and filtered (0.45μM). Virus was extracted using Lenti-X™ concentrator (Clontech Takara) according to the manufacturer’s instructions. The virus containing pellet was resuspended in serum-free neurobasal medium and target cells were infected in the presence of 1μg/ml polybrene (Sigma Aldrich). Two days following infection, cells were selected by growth in puromycin (E2 GSC: 1μg/ml, G7 GSC: 0.5μg/ml, Sigma) containing medium. Drug resistant GSC were further sorted by FACS to isolate the high signal expressing population of iRFP-labelled GSC.

### Cell proliferation assay

GSC were seeded in three wells of a 6-well plate at a density of 5×4^4^ cells per well. Cell number for each well was assessed using a haemocytometer 1, 3 and 7 days after seeding. A median was calculated for each time point. Cell count was normalised to day 1 (n=3 independent repeats per cell line).

### Neurosphere formation assay

GSC were seeded in uncoated 96-well plates at a density of 10 cells per well. Medium was refreshed every week. Spheres were left to grow for 14 days (G7 GSC) or 21 days (E2 GSC) before manual scoring of the 60 inner wells (n=3 independent repeats per cell line).

### Immunoblotting and antibodies

GSC were initially lysed and collected in RIPA buffer (50mM Tris-HCl pH 7.5, 150 mM NaCl, 1 mM EDTA, 1% NP-40), supplemented with complete protease inhibitor (Roche) and PhosSTOP (Roche). Pierce BCA protein assay kit (Thermo Fisher Scientific) was used to determine protein concentration. Western blotting was performed by electrophoresis using 4-12% NuPage Bis-Tris protein gels (Thermo Fisher Scientific) followed by transfer to nitrocellulose membranes. After blocking in 5% non-fat, dry milk membranes were probed with primary antibody (1:1000 unless otherwise stated; Sox2 (Abcam #92494), Nestin (Abcam #22035), Olig2 (R&D Systems #AF2418), GFAP (Santa Cruz #SC-6170) at 4°C overnight. All membranes were incubated with α-tubulin serving as loading control (Sigma #T5168, 1:5000) for one hour at room temperature. Membranes were incubated with Li-Cor secondary antibodies (IRDye 680RD donkey anti-mouse, IRDye 800CW donkey anti-rabbit, IRDye 800CW donkey anti-goat) for one hour at room temperature. Blots were imaged using Li-Cor Odyssey CLx (Li-Cor), acquired and processed using Image-Studio software (Li-Cor) and subsequently arranged using Adobe Illustrator.

### Intracranial Xenografts

All mouse experiments were carried out in accordance with the Animals Act 1986 (Scientific Procedures on living animals) and the regulatory guidelines of the EU Directive 2010 under project licence PPL P4A277133 and ethical review (University of Glasgow). 7-week old, female CD1-nude mice (Charles River, UK) were orthotopically injected with 1×5^5^ iRFP-labelled G7 GSC or E2 GSC into the right striatum. Mice were monitored for the duration of the experiment and humanely sacrificed when they showed neurological (hemiparesis, paraplegia) or general symptoms (hunched posture, reduced mobility, and/or weight loss >20%). To examine intracranial tumour growth, mice were monitored by PEARL imaging (Li-Cor) acquired and processed using Image-Studio software (Li-Cor). For imaging, mice were anaesthetised using isoflurane. Four weeks after tumour injection, a baseline scan was performed. Regular imaging was commenced six weeks (G7 GSC) or eight weeks (E2 GSC) after tumour injection. The percentage change in the iRFP signal as a function of baseline scan was calculated according to the following equation:

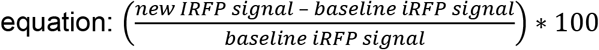

H&E staining and immunohistochemistry (IHC) for Ki67 were performed on formalin-fixed, paraffin embedded brain sections (4μm thick slices). Scanning and image analysis of both was conducted using Leica SlidePath and Halo (Indica Labs). Algorithms were optimised for each stain individually and automated, quantitative analysis undertaken with Halo software (Indica Labs).

### Statistical analyses

All statistical analyses were executed with Prism software version 8.4.2 (GraphPad, La Jolla, CA, USA). Significance of neurosphere formation assay was calculated using Welch’s t-test. For analysis of correlation of Ki67 positivity and iRFP signal increase, single linear regression and Spearman’s *r* were used.

## Results

### Generation of iRFP-expressing glioblastoma stem-like cells

Near-infrared fluorescent proteins (iRFP) with excitation and emission in the 700 and 800 nm spectra allow tumour detection in deep tissues with only minimal interference from tissue autofluorescence ^11^. Since the use of iRFP in mouse models was first described in 2011, it has been shown to be a powerful and reliable tool to measure cell proliferation *in vitro* and *in vivo* ^11,14^. We deployed a retroviral transduction approach to express iRFP in E2 and G7 GSC, two primary patient-derived cell lines originally derived from fresh patient tissue as previously described ^12,13^. We tailored the protocol to reduce the number of manipulations so as to minimise effects on the heterogeneous nature of the GSC cultures. To avoid culturing GSC in FCS containing medium, which would drive cell differentiation, the iRFP-encoding retroviruses were resuspended in neurobasal medium. 48 hours after infection, enrichment of iRFP-expressing GSC was commenced using puromycin selection, followed by fluorescence activated cell sorting (FACS) for a cell population with higher signal intensity (**Figure 1a**). Due to the infiltrative nature of E2 GSC xenograft growth, stricter gating parameters were applied to ensure subsequent detectability of the iRFP signal *in vivo*. There were no detectable differences between the morphology and proliferation rates of iRFP-expressing GSC and their matching parental controls (**Figure 1b, c**).

**Figure 1.**
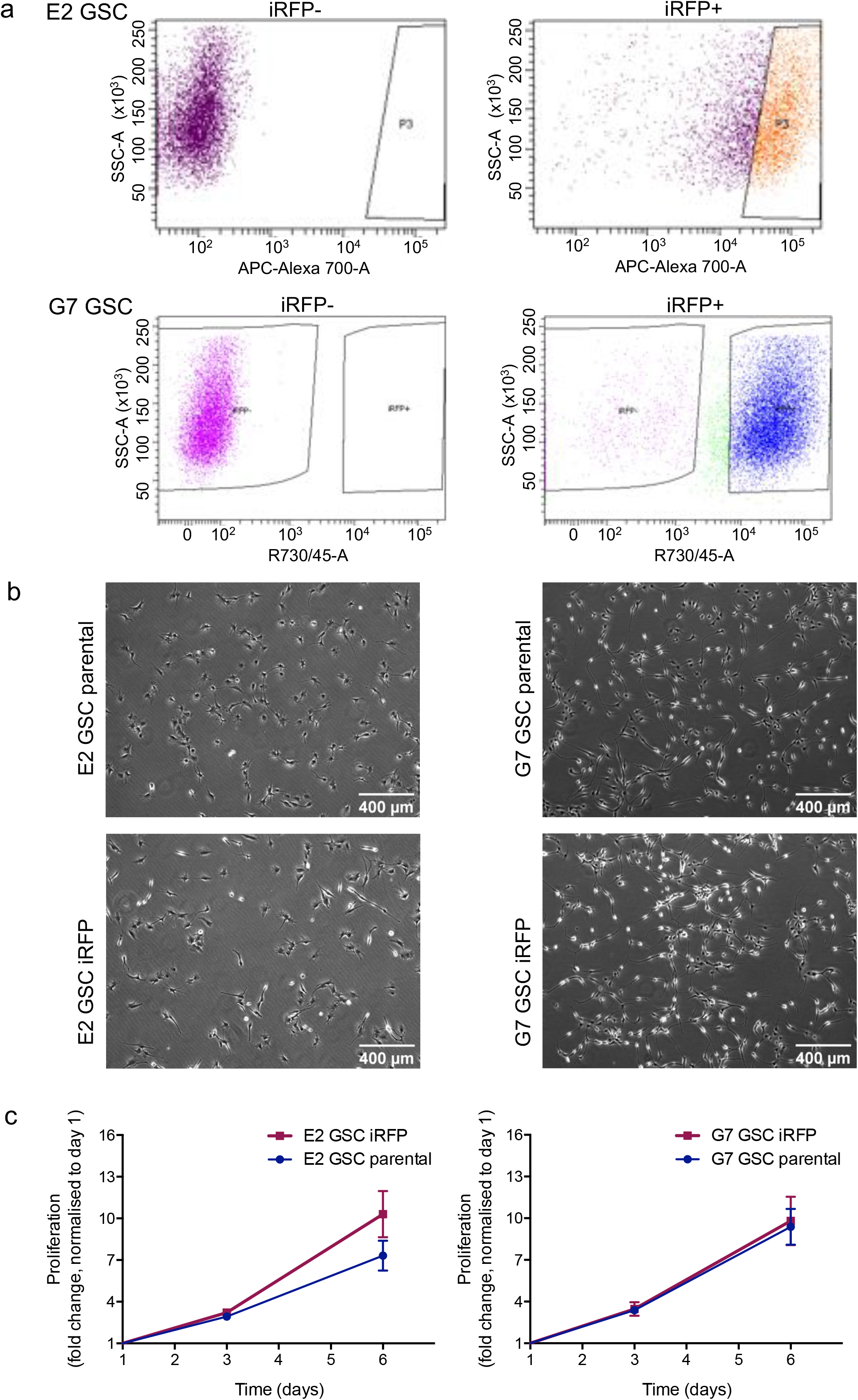
Generation of iRFP-expressing glioblastoma stem-like cells. (a) Flow cytometric plots showing sorting for higher signal iRFP-overexpressing E2 and G7 GSC (upper panel: P3 = high signal iRFP positive cells; lower panel: iRFP+ = iRFP positive cells) (b) Representative images of monolayer E2 and G7 GSC parental (upper panel) and iRFP-expressing cells (lower panel), respectively. (c) Proliferation assay of E2 and G7 GSC counted at the indicated times after plating. Cell count was normalised to day 1. Error bars represent mean +/−SD from n=3 independent experiments.

### iRFP-expressing GSC retain stem-like features

Cancer stem cells are understood to be the tumourigenic drivers of glioblastoma^15–17^. Consequently, extensive research has been devoted to refining glioblastoma culturing methods so as to preserve the specific stem-like properties of patient tumour specimens ^9,12,15,18^. The microenvironment, including medium composition and culture conditions, inevitably affect the characteristics of stem cell populations ^19^. Their self-renewal capacity not only encourages rapid adaption to alterations in the microenvironment, but can also drive resistance to cancer therapies ^20^. Thus, we were keen to determine whether changes in stem cell characteristics had occurred after retroviral transduction. The capacity to form neurospheres (as a proxy for tumour initiation) was conserved in the iRFP-expressing GSC, although a slight (but not statistically significant) decrease of neurosphere forming capacity was detectable in the E2 GSC iRFP (**Figure 2a**). Furthermore, the expression of cell lineage-specific neural stem cell (SOX2, Nestin, OLIG2) and astrocytic differentiation (GFAP) markers was maintained upon retroviral transduction and expression of iRFP, although the Olig2 and GFAP protein levels in E2 GSC were moderately decreased (**Figure 2b**). This decrease is likely to be caused by the more stringent FACS criteria applied to these cells (see **Figure 1a**), further highlighting the sensitivity of patient-derived primary glioblastoma cell lines to manipulation. Taken together, these data demonstrate that our approach to achieve iRFP expression is unlikely to influence GSC stem-like behaviour.

**Figure 2.**
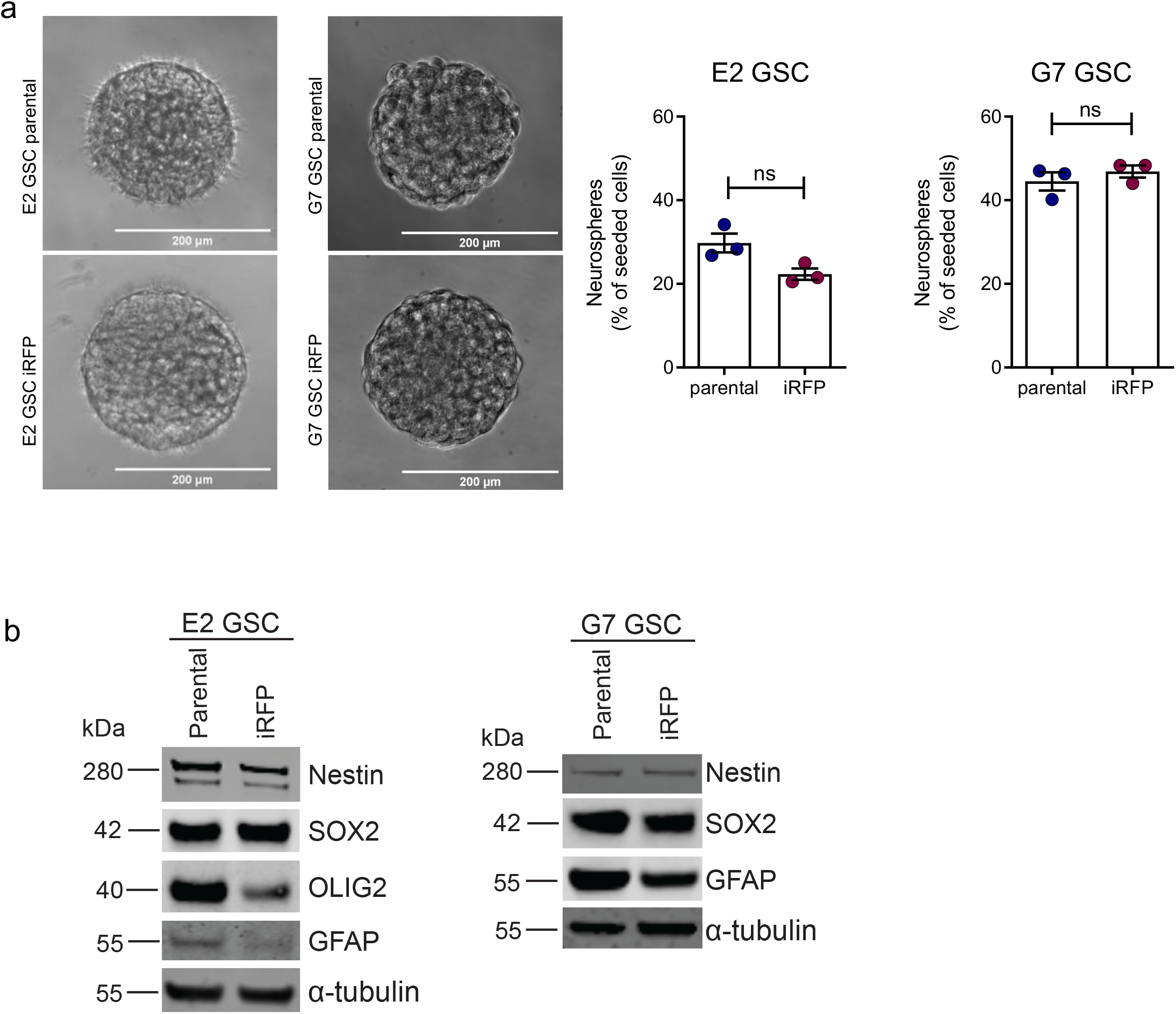
iRFP-expressing GSC retain stem-like features. (a) Representative images of neurosphere growth from E2 and G7 GSC parental (upper panel) and iRFP-expressing cells (lower panel), respectively. Quantification of neurosphere formation capacity by E2 parental vs. iRFP GSC (p=0.059) and G7 parental vs. iRFP GSC (p=0.419). Error bars represent mean +/−SEM from n=3 independent experiments; ns, nonsignificant. (b) Immunoblotting for cell-line specific neural stem cell markers (SOX2, OLIG2, Nestin) and astrocyte lineage differentiation marker GFAP in E2 and G7 GSC parental compared with paired iRFP GSC, respectively. Alpha-tubulin served as loading control. Representative loading controls given (E2 GSC: membrane for SOX2/Nestin, G7 GSC: membrane for GFAP). All loading controls are displayed with full-length blots in Supplementary Figure 1.

### Quantitative imaging of iRFP-expressing glioblastoma in vivo

We next determined whether iRFP was suitable for measuring glioblastoma growth *in vivo*. For this purpose, we generated orthotopic xenografts using E2 GSC iRFP and G7 GSC iRFP. When implanted into the right striatum of 7-week old CD1-nude mice, both iRFP-expressing cell lines retained the growth patterns characteristic of their parental counterparts. While iRFP-expressing G7 GSC grew mainly as a solid tumour with invasive margins, E2 GSC iRFP disseminated throughout the brain as widely scattered tumour cells (formerly described as gliomatosis cerebri) to such an extent that the presence of a solid tumour at the injection site was not readily discernible (**Figure 3a**). To date, no imaging tool has been capable of reliably detecting or quantitatively monitoring growth of highly invasive tumour xenografts, such as those engendered by E2 GSC. To determine whether iRFP expression enabled bona fide quantification of these highly disseminated tumours, we implanted 1×5^5^ iRFP-expressing E2 GSC into nude mice as before. Mice were initially scanned using a PEARL imager to establish a baseline, and this was performed 4 weeks after surgery to allow for sufficient recovery and wound healing. Images from single mice were acquired within 30 seconds, followed by a short awakening and recovery phase. Based on previous studies using E2 GSC, we began imaging the mice at regular intervals from week 8 until clinical endpoint or 31 weeks after surgery ^21,22^. Analysis of the signal intensity over time revealed that a measurable increase in signal (with reference to the 4 week baseline) was detected as early as 16 weeks after surgery, delineating individual rates of signal increase for each animal (**Figure 3b, c**). To investigate the utility of iRFP imaging in PDX models growing as solid tumours with invasive margins, we used orthotopic G7 iRFP GSC xenografts. In this experiment, imaging commenced six weeks after tumour cell injection. Individual signal increase patterns for each of the three mice were detectable from 8 weeks following implantation, thus indicating that iRFP-expressing GSC may be used to monitor the growth of glioblastoma PDXs over an extended time course (**Figure 3d, e**).

**Figure 3.**
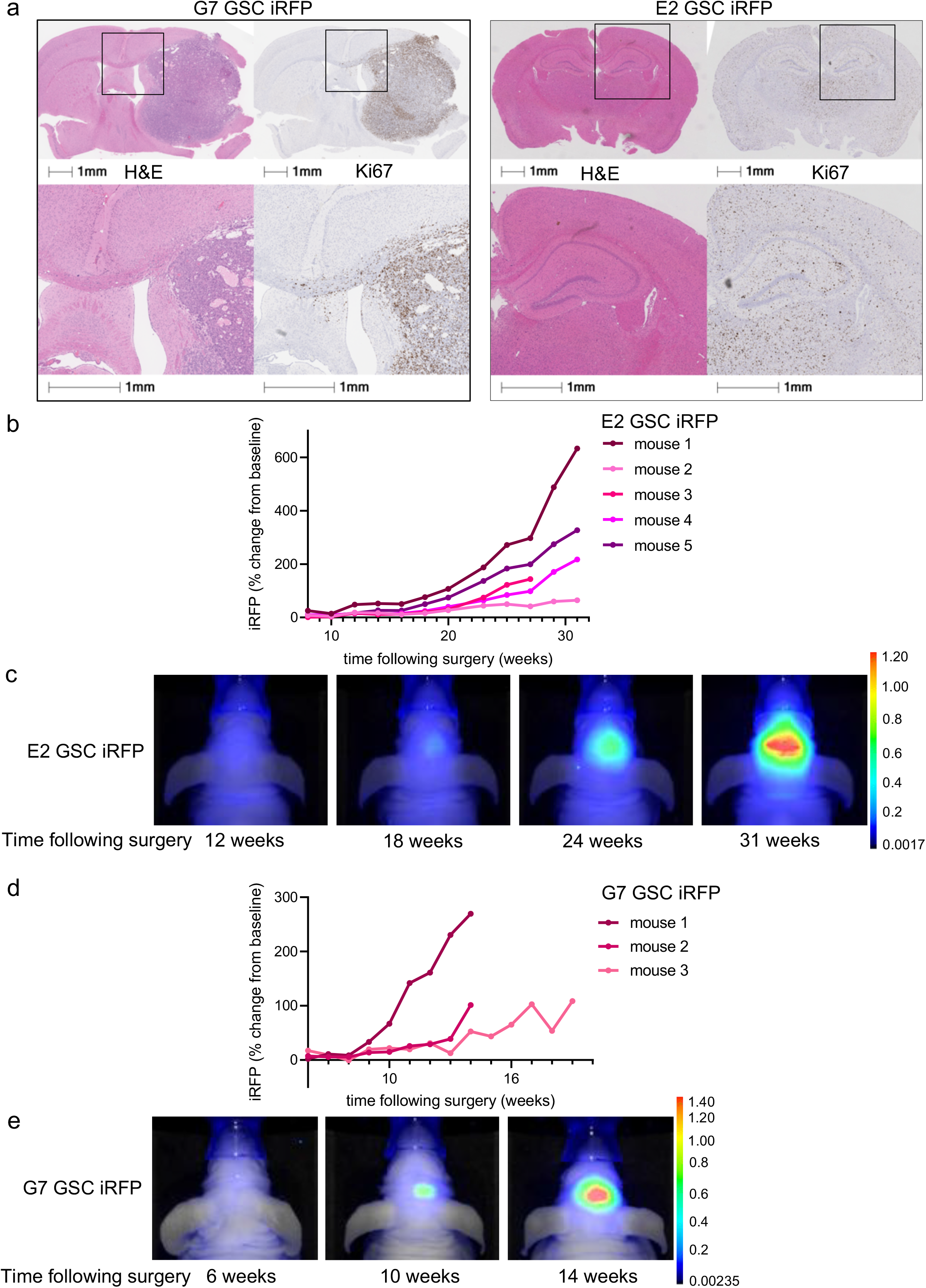
Quantitative imaging of iRFP-expressing glioblastoma *in vivo*. (a) Representative images of haematoxylin and eosin (H&E) and Ki67 immunohistochemistry of brain sections containing orthotopic tumours derived from intracranially injected G7 or E2 GSC iRFP. (b) E2 GSC iRFP were injected intracranially into five mice. iRFP signal increase was measured at indicated times (c, e). Pseudocolour representations of iRFP signal increase over time in individual mice bearing E2 or G7 GSC iRFP tumours, respectively, as detected by PEARL scans (700nm channel) (d) G7 GSC iRFP were injected intracranially in three mice. iRFP signal increase measured at indicated times.

### iRFP signal correlates with histological tumour burden

To determine whether the fluorescent signal emanating from iRFP-expressing GSC tumours correlated with histological tumour burden, we *ex vivo* imaged brains bearing G7 GSC iRFP xenografts. We then fixed these brains and stained them using H&E, thus allowing comparison of the fluorescence and histological fields. We found that in mice implanted with G7 GSC, iRFP fluorescence faithfully delineated the extent of intraparenchymal spread that had been detected using iRFP imaging. Strikingly, imaging was sufficiently sensitive to illustrate the central core necrosis that was evident following H&E staining (**Figure 4a**). Since G7 iRFP tumours grew approximately spherically *in vivo*, we extrapolated tumour volume from the mean diameter in H&E and correlated it with the absolute iRFP signal measured at the final imaging obtained before endpoint. Notably, the fluorescent signal emanating from iRFP-expressing G7 GSC tumour bearing mice was highly concordant with the calculated tumour volume (**Figure 4b**). As highly invasive models such as E2 GSC do not permit measurement of a defined tumour bulk, we sought to examine the accuracy with which iRFP signal intensity represented tumour burden. Using the proportion of Ki67-positive cells across the brain section as a proxy, we observed that the burden of disseminated E2 GSC tumour cells varied substantially between animals, similar to the intravital imaging data (**Figure 4c**). Interestingly, we found that the relative iRFP signal increase at endpoint correlated strongly with the relative prevalence of Ki67-positive cells as quantified by automated image processing software (Spearman correlation *r*=0.9). Taking into account the limited number of animals, the observed trend towards statistical significance (p=0.0755) supports the reliability of the model (**Figure 4d**). Taken together, our data demonstrate that near-infrared labelling of highly invasive primary patient-derived glioblastoma cells is a powerful tool to quantitatively monitor their growth *in vivo*.

**Figure 4.**
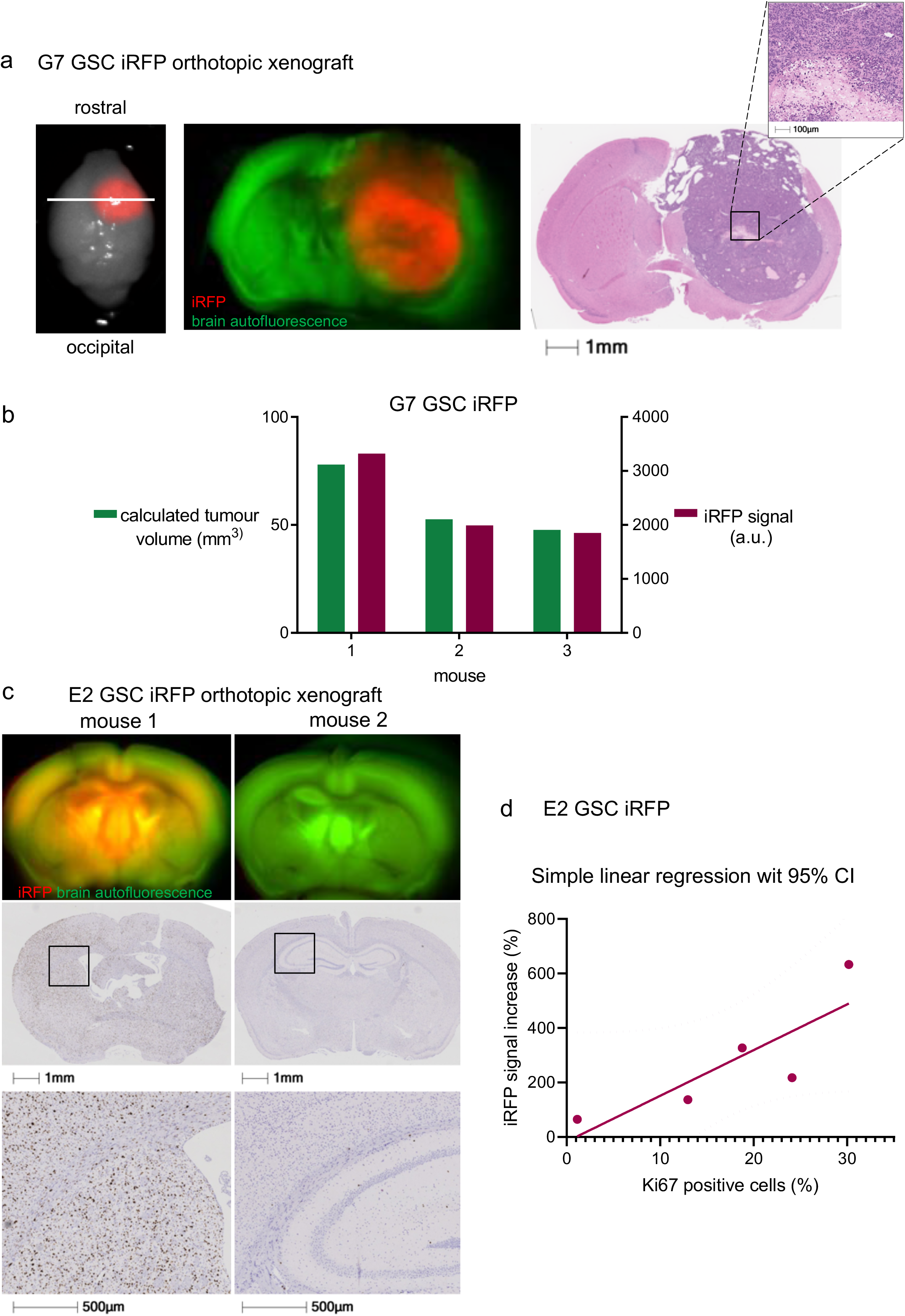
iRFP signal correlates with histological tumour burden. (a) Representative *ex vivo* PEARL and Odyssey scans of G7 GSC iRFP tumour in a mouse brain after resection (left), and on coronar section (middle, denoted by white line in left image) in comparison to matching H&E stain (right). 700nm scan (red) was used to visualise the tumour, brain autofluorescence was detected in 800nm channel (green). (b) G7 GSC iRFP signal detected with PEARL imager (a.u., arbitrary units) corresponds with calculated spherical tumour volume (mm^3^), extrapolated from mean diameter in H&E stains. (c) Upper panel: Odyssey scans of coronar sections from E2 GSC iRFP bearing mice. 700nm scan (red) was used to visualise tumours, brain autofluorescence was detected in 800nm channel (green). Middle and lower panel: corresponding Ki67 staining and magnification. (d) Simple linear regression correlating E2 GSC iRFP signal with Ki67 positive tumour cells in five mice, quantified with Halo analysis software (p=0.0755, Spearman correlation *r*=0.9).

## Discussion

Reporter gene-based technologies, such as luciferase-based bioluminescence, are widely used but have particular limitations especially with regard to quantitative longitudinal assessment of tumour xenograft growth. In this paper we show that the use of cell lines stably expressing iRFP overcomes many of these limitations. We have demonstrated that it is feasible to manipulate primary glioblastoma stem-like cells by stably overexpressing the near-infrared fluorescent protein iRFP without overtly affecting their particular stem-like properties. Signal intensity is consistent and reliable over time and permits monitoring of orthotopic tumour growth *in vivo* with a high degree of accuracy. This is especially valuable for highly invasive cell lines such as E2 GSC, which cannot be detected or quantified with conventional imaging methods such as MRI. As iRFP enables cost-effective, repeated and comparatively high throughput non-invasive imaging of mice bearing orthotopic glioblastoma PDX, it could be used to determine individual treatment start dates for mice when they reach a certain threshold, mimicking more closely the clinical scenario. Additionally, this method may also be used to stratify mice into cohorts depending on their actual biological tumour burden. This application serves to reduce the number of mice needed for statistical power in heterogenous primary glioblastoma PDX-models. A further advantage of iRFP expressing tumours is the ability to FACS-sort tumour cells without the need for antibody labelling, enhancing cell viability and leaving cell surface molecules unperturbed.

In summary, our results show that iRFP serves as a reliable and sensitive tool to monitor intracranial tumour growth in both highly invasive and more localised primary glioblastoma orthotopic xenograft models without affecting their phenotypes. This new technique provides significant benefits for the design, execution and analysis of *in vivo* studies using glioblastoma PDX.

## ACKNOWLEDGEMENTS

We thank Professor Colin Watts for providing the E2 and G7 GSC, Dr. Andreas Hock for providing the pBABE iRFP IRES puro construct and Dr. David Bryant for providing the HEK293-FT cells. In addition, we thank Tom Gilbey, Colin Nixon, the Histology service, Biological Services, and all core services at the CRUK Beatson Institute for their invaluable assistance. Many thanks to Dr. Matthew Neilson for his invaluable support and advice on statistical analysis, and Dr. Florian Bock and Dr. Catherine Winchester for critical reading and assistance in the preparation of this manuscript.

This work was supported by CRUK core funding to the Beatson Institute (A17196) and to J.C.N. (A18277), a CRUK Programme Foundation Award (C40872/A20145) to S.W.T., CRUK Clinical Research Fellowship to A.L.K. (A23220), funding by the University of Glasgow to D.K. and A.C, and funding by the Beatson Cancer Charity and Cancer Research UK RadNet Centre Glasgow to K.S..

## AUTHOR CONTRIBUTIONS

A.L.K. and D.K. performed experiments. K.S., C.C. and L.M. provided assistance with the intracranial xenograft model and related imaging. A.L.K. and D.K. designed experiments and analysed the data. A.L.K., D.K., S.W.T. and J.C.N. wrote the manuscript. A.C., S.W.T. and J.N. directed the study.

---

**Supplementary Figure 1.**
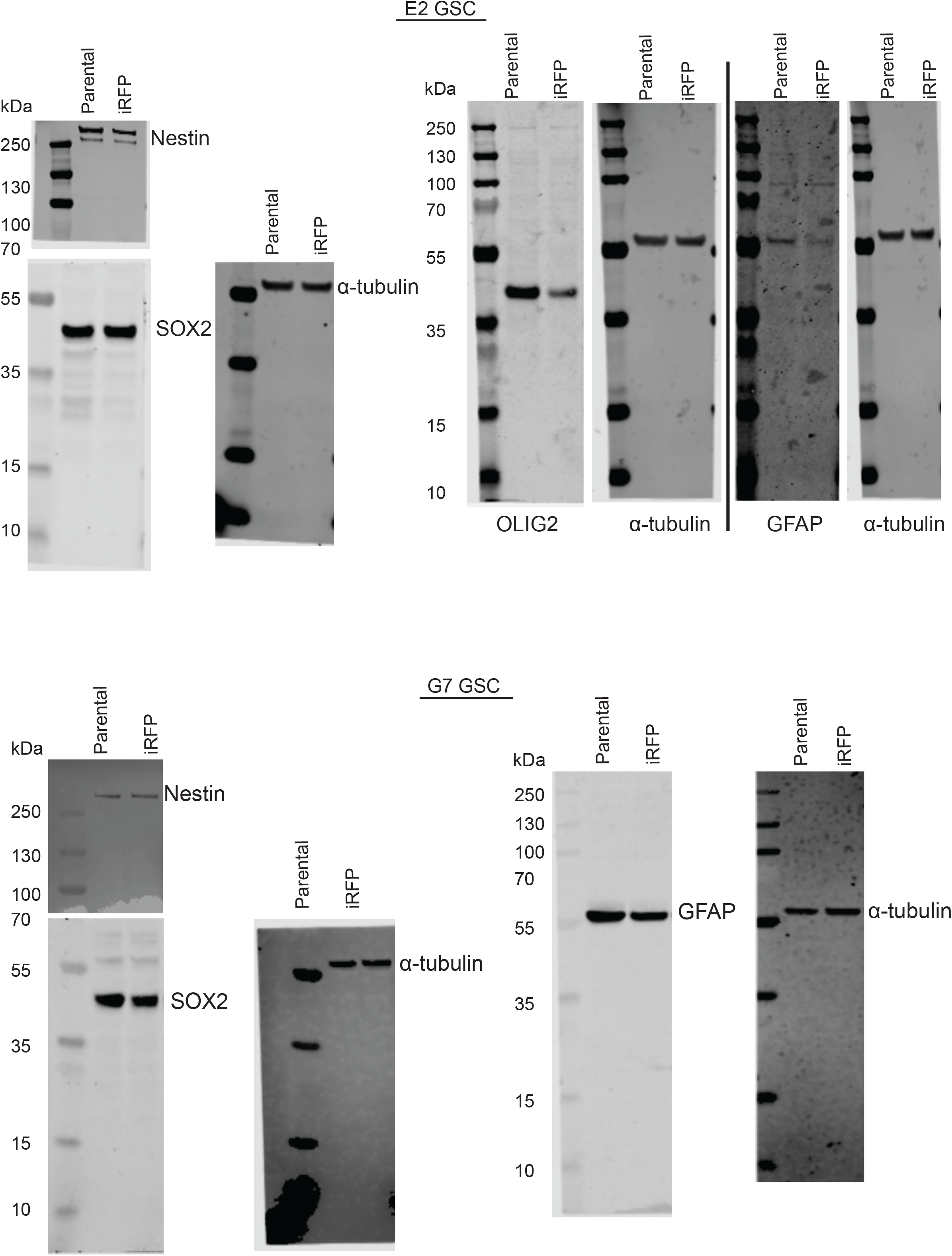

